# Linking functional traits with tree growth and forest productivity in *Quercus ilex* forests along a climatic gradient

**DOI:** 10.1101/2020.12.12.422386

**Authors:** Pablo Salazar Zarzosa, Aurelio Diaz Herraiz, Manuel Olmo, Paloma Ruiz-Benito, Vidal Barrón, Cristina C. Bastias, Enrique G. de la Riva, Rafael Villar

## Abstract

Plant functional traits are highly adaptable to changes in climatic factors and nutrient availability. However, the intraspecific plant response to abiotic factors and the overall effect on plant growth and productivity is still under debate. We studied forest productivity for 30 *Quercus ilex* subsp *ballota* forests in Spain along a broad climatic gradient of aridity (mean annual precipitation from 321 to 1582 mm). We used linear mixed models to quantify the effect of climatic and edaphic factors on functional traits, and to study the effect of functional traits and abiotic factors on the relative growth rate (RGR) of adult trees. Then we used piecewise structural equation models (SEMs) to determine the causal effect of intrinsic and extrinsic factors on forest productivity. Our results showed that forest productivity is positively affected by forest biomass and RGR, which are mainly affected by functional traits and tree biomass, respectively. In conclusion, intraspecific variability of functional traits have a significant effect on plant biomass and growth, which ultimately explain forest productivity in *Quercus ilex*.

## Introduction

Functional traits are biological attributes that directly or indirectly affect plant fitness and generally reflect plant adaptation to local environmental conditions (Lavorel et al., 1997; Violle et al., 2007). Changes in these traits are made to minimize their building costs and maximize functional efficiency. For instance, drought avoidance creates a trade-off between water conservation and nutrient acquisition. Wood density and leaf mass per area (LMA) increase in arid environments to reduce the risk of cavitation and increase defense functions at the expense of lower plant biomass and growth (Chave et al., 2009; Vilà-Cabrera et al., 2015). Understanding how functional traits vary across environmental gradients is critical to determine plant functioning and their ecological strategies in contrasting environmental conditions (Westoby, 1998), especially relevant under the ongoing climatic change scenario.

Empirical studies have shown that functional traits covariate with abiotic variables with a direct effect on the relative growth rate (RGR; Chave et al., 2009; Salgado-Luarte and Gianoli, 2017; Violle et al., 2007). At the local scale, the effects of functional traits have an important effect on RGR (Chaturvedi et al., 2011; Dong et al., 2020). Whereas at larger spatial scales tree size is used as the main plant trait and abiotic conditions are summarized through temperature or precipitation to explain plant growth (Enrique G de la Riva et al., 2016; Moore et al., 2020). In fact, a weak relationship between LMA and RGR has also been found when plant size is taken into account (Gibert et al., 2016; Wright et al., 2010), arguing that LMA is not a clear physiological trait and should be replaced by mechanistic traits (like leaf nutrients or wood density) more associated with plant fitness (Rosas et al., 2019). The direct and indirect effect of functional traits on RGR is key to understand how plants adapt to environmental changes, and whether or not this has any effect on ecosystem functioning.

Additionally, variation in RGR is associated with intrinsic factors as the tree biomass and abiotic factors such as soil nutrient availability and climate (Antúnez et al., 2001; Cornelissen and Thompson, 1997; Lambers et al., 2008). However, it is still uncertain whether these factors have a direct or indirect effect on plant growth. For instance, Bu et al. (2019) found that changes in abiotic conditions did not directly affect plant growth, but indirectly via changes in plant functional traits. Similarly, it has been shown that RGR negatively correlates with tree biomass during the stand development; however, this relationship disappears when the trees reach the maturity stage (Ruiz-Benito et al., 2015). Therefore, the source of variation in RGR is currently a growing field of knowledge.

Several large-scale studies have analyzed how changes in functional traits between species determine growth responses and forest demography (Kunstler et al., 2016; Ruiz-Benito et al., 2017). It is not well known however the role of intraspecific variability in determining forest responses (Wang and Hamzah, 2018), although it has been suggested that intraspecific variation determines species distribution under a warming climate (Valladares et al., 2015). Furthermore, it has been shown that intraspecific variability can be larger than interspecific variability (Fajardo and Siefert, 2016), and this variation influences the interactions among and between organisms that ultimately drive the functional community (Bastias et al., 2017; Siefert et al., 2015). Therefore, understanding the source of intraspecific functional trait variations, as well as its contribution to plant growth and ecosystem function is critical to understand forest responses under changing conditions. We focus on the intraspecific variability of *Quercus ilex* (subsp ballota) dominated forests, a key tree species in Mediterranean forests characterized by high plasticity and resistance to drought (Aranda et al., 2004; Quero et al., 2006).

Our goal is to assess the relationships between local-scale variables (plant functional traits and abiotic conditions) and large spatial and temporal ecosystem functions as annual forest productivity and RGR (Fig. 1). We hypothesized that the annual forest productivity (calculated as kg ha^-1^ year^-1^) is related to the forest biomass (in kg ha^-1^) and RGR (kg kg^-1^ year^-1^). The former can be decomposed in the mean tree biomass and the tree density, whereas the latter can be explained by functional traits and abiotic factors. To study this, we combined data from the Spanish National Forest Inventory (NFI) with field sampling to include functional trait data and soil characteristics. Our specific objectives are: (1) to determine the functional traits that are most sensitive to abiotic factors (climate and soil properties); (2) to study the effect of abiotic factors, functional traits, and forest structure (tree biomass and density) on relative growth rate under natural conditions; and (3) to model forest productivity as a result of these ecological, physiological and environmental factors.

**Figure 1.**
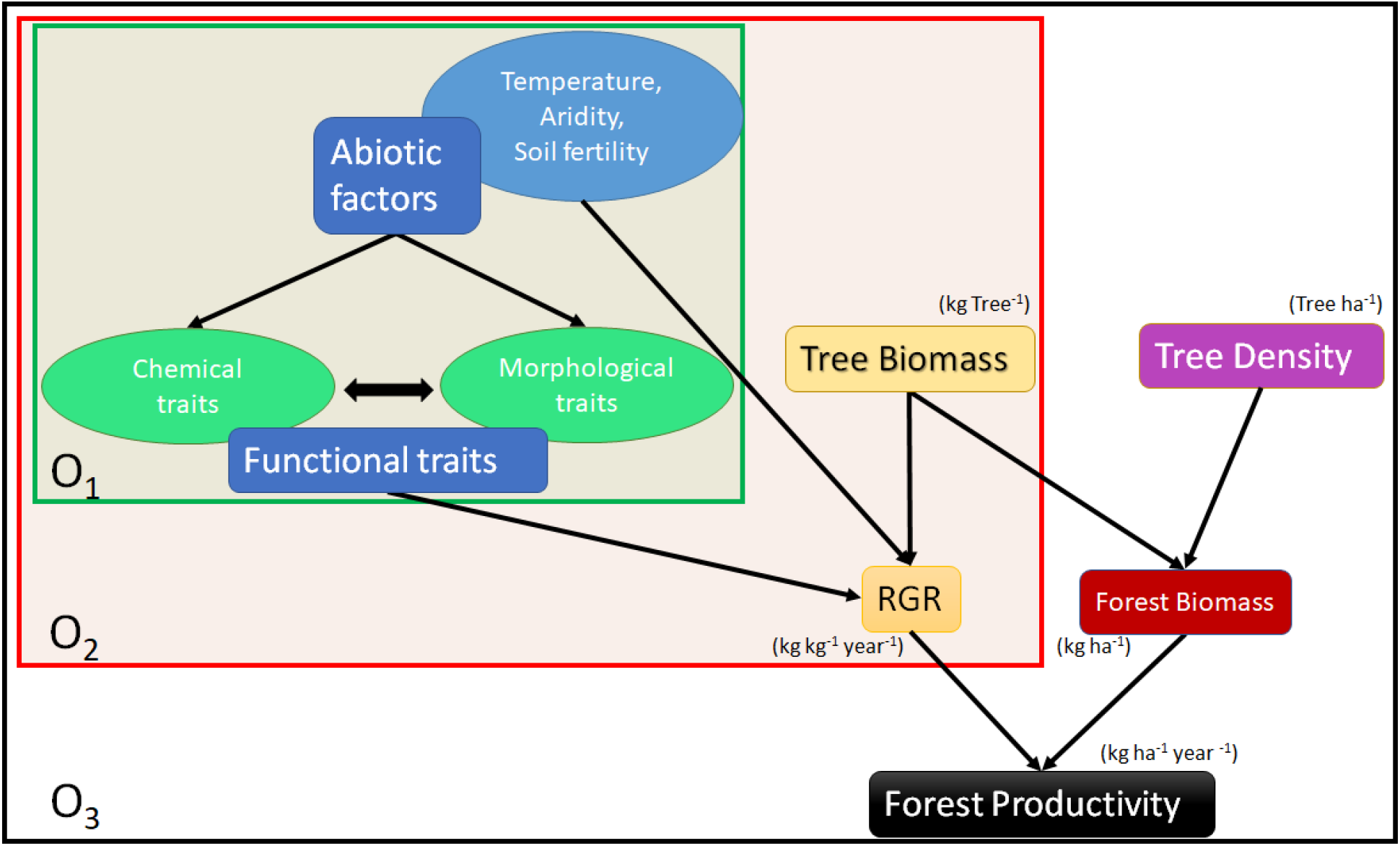
Theoretical model explaining the hypothesized effects of abiotic factors on functional traits (Objective 1, O_1_). The effect of functional traits, abiotic factors and tree biomass (kg tree^-1^) on relative growth rate (RGR, kg kg^-1^ year^-1^; Objective 2, O_2_), and its effect on forest productivity (kg ha^-1^ year^-1^) and forest biomass (Objective 3, O_3_).

## Material and methods

### Study species

We studied *Quercus ilex* subsp *ballota* (hereafter *Q. ilex*), an evergreen-sclerophyllous and drought-resistant species, which is extensively distributed throughout the Mediterranean basin (Caudullo et al., 2017; Quero et al., 2011; Fig. 2). This species is shade-tolerant characterized by an anisohydric behavior (Sade et al., 2012). It is deep-rooted and has a great capacity to maintain high stomatal conductance during long dry periods (Barbero et al., 1992; Quero et al., 2011).

**Figure 2.**
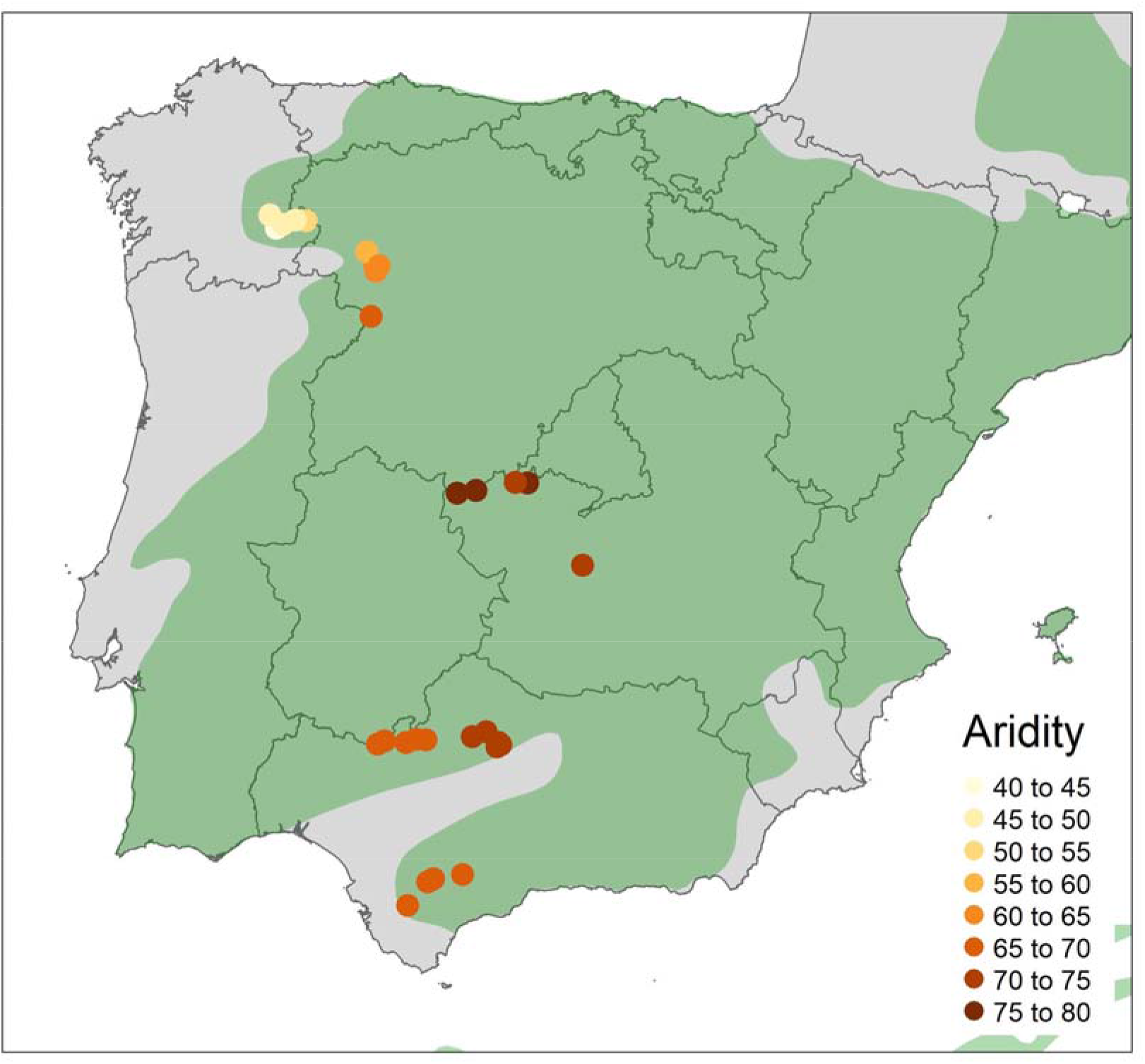
Spatial distribution of the 30 sampled plots dominated by *Quercus ilex* across an environmental gradient in Spain. Dot color indicates average aridity index.

### Spanish Forest Inventory and experimental design

The Spanish Forest Inventory is a nationwide program that establishes plots in forested areas of Spain each km^2^. We selected thirty plots from the inventory dominated by *Q. ilex*. The plots were selected covering six contrasting climatic regions according to Köppen-Geiger Climate Classification. The mean annual temperature oscillates between 10.9 to 17.3 °C and the annual precipitation oscillates from 321 to 1582 mm. More details about the selected plots can be found in Table 1.

**Table 1.** Descriptive statistics of the functional traits, forest traits, and abiotic factors measured in this study.

| Abr. | variable | mean | sd | min | max | CV |
| --- | --- | --- | --- | --- | --- | --- |
| LA | Leaf area (cm<sup>2</sup>) | 4.45 | 1.64 | 1.66 | 10.61 | 36.93 |
| LD | leaf density (g cm<sup>-3</sup>) | 0.54 | 0.07 | 0.29 | 0.82 | 13.73 |
| LMA | Leaf mass per area (g m<sup>-2</sup>) | 215.5 | 35.8 | 114.9 | 321.0 | 16.60 |
| LT | Leaf thickness (μm) | 402.9 | 57.0 | 233.2 | 558.8 | 14.16 |
| SWD | Stem-wood density (g m<sup>-3</sup>) | 1.14 | 0.05 | 0.88 | 1.35 | 4.39 |
| SWDMC | Stem-wood dry matter content (%) | 61.41 | 4.93 | 51.93 | 85.69 | 8.03 |
| TB | Tree biomass (kg tree<sup>-1</sup>) | 518 | 702 | 16 | 4623 | 136 |
| TD | Tree density (N° tree ha<sup>-1</sup>) | 1097 | 2340 | 32 | 12732 | 213 |
| FB | Forest biomass (Mg ha<sup>-1</sup>) | 26.97 | 10.82 | 8.65 | 53.21 | 40.12 |
| FP | Forest productivity (Mg ha<sup>-1</sup> year<sup>-1</sup>) | 1.91 | 1.38 | 0.40 | 5.43 | 72.17 |
| RGR | Relative growth rate ( g kg<sup>-1</sup> year<sup>-1</sup>) | 50.28 | 46.13 | 0.45 | 236.37 | 91.74 |
|  | Clay (%) | 10.01 | 9.83 | 1.04 | 44.13 | 98.15 |
| MAP | Mean annual precipitation (mm) | 634.8 | 145.5 | 358 | 878 | 22.92 |
| MAT | Mean annual temperature (°C) | 14.31 | 1.82 | 10.90 | 17.30 | 12.75 |
| AI | Aridity index | 65.22 | 9.70 | 42.63 | 79.37 | 14.87 |

We used the diameter at breast height (d.b.h.) data from the third National Forest Inventory (1997-2007) available for the entire country. Tree biomass (kg tree^-1^) was calculated from d.b.h. with the allometric equation in Ruiz-Peinado et al. (2011). The number of trees within the plot was used to calculate the tree density (No. trees ha-1), and forest biomass (Mg ha-1) was calculated as the sum of the biomass of all trees within the plot divided by the plot area. In 2018, we re-measured the diameter at breast height of 5 trees in each selected plot to calculate tree biomass and forest biomass in the last eleven years (2007 - 2018 period). Moreover, we collected leaf and wood material from each tree to measure key functional traits related to plant ecological strategy under water deficit.

Tree relative growth rate (RGR) was calculated as: RGR = [Ln (B2) - Ln (B1)] / [t2 – t1], where B is tree biomass and t is time, with 2 and 1 referring to our sampling survey data (i.e. 2018) and NFI3 data (i.e. 2007), respectively. Forest productivity (kg ha-1 year-1) was calculated as the difference of forest biomass between our sampling data and NFI3 for trees alive in both consecutive inventories divided by the time difference between surveys (between 9 and 11 years).

### Functional traits measurements

Leaf and wood material collection was done in late spring and summer, between May and September 2018, when the leaf formation is completed. In each plot, five *Q. ilex* trees were sampled taking two branches per individual. The samples were conservated in a damp paper in a portable refrigerator and transported to the laboratory. Fully mature one-year-old leaves were taken from each branch and processed independently. For a subsample of five leaves, we scanned them (excluding petiole) and the leaf area (LA) was analyzed using Image Pro software (Media Cybernetics, MD, USA). Then, we measured the leaf thickness (LT) using a micrometer (Electronic Digital Micrometer Comecta, Barcelona, Spain). After that, the leaves were dried on a stove (60°C for two days) to calculate leaf dry mass. Leaf mass area (LMA, g m^-2^) was calculated as dry mass (g) / leaf area (m^2^), leaf density (LD; g cm^-3^) as LMA (g m^-2^) / thickness (µm) and leaf dry matter content (LDMC; g g^-1^) = dry mass (g)/ fresh mass (g). Another leaf subsample was dried on a stove (60 °C for 2 days) and reserved for nutrient analysis.

At the moment of nutrient analyses, leaves were dried at 70°C for 24 h before grinding with a stainless steel grinder. The leaf N concentration was measured using an elemental analyzer (Eurovector EA3000), and we also measured the leaf concentrations of macronutrients (P, K, Ca, and Mg) and micronutrients (Na, Fe, Mn, Cu, and Zn). To do so, leaf samples were digested using a mixture of nitric acid and perchloric acid. Phosphorus concentration was determined according to the molybdate blue method (Murphy and Riley, 1962). Ca and Mg were determined by atomic absorption spectrophotometry, while K and Na were determined by atomic emission spectrophotometry. Lastly, Fe, Mn, Cu, and Zn were determined by atomic absorption spectrophotometry.

Additionally, two stem samples of about 5 cm length and 1 cm diameter were collected from the two branches of each individual. We weighed them to calculate the stem-wood fresh mass (in g) and we calculated the stem volume using Archimedes′ principle (displaced volume of water). Stems were dried at 60°C for 4 days to obtain stem-wood dry mass (SWDM). Stem-wood density (SWD, g cm^-3^) was calculated as SWDM/ wood volume, and stem-wood dry matter content (SWDMC) was calculated as stem-wood dry mass/ stem-wood fresh mass. All measured functional traits were calculated following the methodology described in Pérez-Harguindeguy et al. (2013).

### Abiotic factors

In each plot, four random points were selected and soil samples were taken at 0-20 cm depth. The soil cores obtained were mixed in two independent samples. Later, the samples were dried at room temperature, ground, sieve (2 mm), and stored in plastic closed bags. Soil texture was determined in 10 g of soil using the Robinson pipette method (Gee et al., 1986). Soil organic carbon was measured by the Walkley and Black (1934) method with potassium dichromate. After a partial extraction of available soil nutrients, we measured soil macronutrients (P, K, Ca, and Mg) and micronutrients (Na, Fe, Mn, Cu, and Zn). To determine P content, we use the NaHCO_3_ 0.5M extraction (Olsen et al., 1982), and for Na, Ca, K and Mg, soil samples were extracted by 1 M NH_4_OAc at pH 7. Fe, Zn, Mn, and Cu were determined after extraction with a solution containing 0.1 M triethanolamine (TEA), 0.005 M diethylenetriaminepentaacetic acid (DTPA), and 0.01 M CaCl_2_ (Norvell and Lindsay, 1972). Then, soil nutrient analysis was made using the same methodology described before for leaf nutrients.

Moreover, mean annual temperature (MAT) and precipitation data (MAP) were downloaded from the WorldClim database (Hijmans et al., 2005). Additionally, we downloaded the precipitation in the driest month (DMP) and the temperature in the driest month (DMT) in Celsius degrees to calculate the modified Martonne aridity index (J., 1979) as (AI): AI = {[MAP] / [MAT + 10]} + {[12 x DMP] / [DMT+10]/ 2}. However, since high AI indicates high water available, we transformed this variable as follows: new AI = 100 -AI.

### Statistical analyses

To reduce the amount of soil nutrient variables (11 variables), a principal component analysis (PCA) for all soil nutrients (S) was carried out (Supplementary Material Fig. S1). The main axes (S1 and S2) explained 40.9% and 20.7% of the variance, respectively (Fig. S1A, Supplementary material). The axis S1 was positively related to Ca and N concentration, while the axis S2 was positively related to P and Fe concentration (Fig. S1B, Supplementary material). To explore the overall relationships between abiotic factors (soil and climate variables) and avoid collinearity in the models, a correlation matrix with all potential abiotic variables was performed (Figure S2A. Supplementary material). Due to the high correlation between MAT, MAP, and AI, we exclude both MAT and MAP from further analysis.

Similarly, we carried out a PCA for the leaf nutrients (L) to reduce the number of variables (Supplementary Material Figs. S3). In this case, the results were also used to analyze the leaf nutrient variability and the variables that drive the most variation. Furthermore, to explore the overall relationships between functional traits and avoid collinearity in the models, a correlation matrix was made (Figure S2B. Supplementary material). The LMA, LD, LA, and LT showed a tight correlation between them, while SWDMC was negatively correlated with SWD. Therefore, only LMA and SWD were included as explanatory variables.

To quantify the effect of abiotic factors (climate variables, soil texture, and nutrients) on chemical and morphological leaf traits, we carried out linear mixed-effects (LME) models for each response variable at tree level (Objective 1) using standardized data. To explore the effect of abiotic factors (climate variables, soil texture, and nutrients) and functional traits on the relative growth rate, we carried out an LME model at the tree level (Objective 2). In both cases, we nested the plot and the climatic region as random factors to consider the no independence of our dataset, and we used a gaussian family function to fit the models.

Finally, to explain the causal effect of these variables on forest productivity, we built two piecewise structural equation models (SEM) using first tree level data to explain the RGR and, second, plot-level data to explain forest productivity (Objective 3). The structure of the hypothesized causal relationships between the selected variables was set based on our expected relationships in Fig. 1. As forest productivity and forest biomass describe the tree biomass of the whole plot, we decided to split the analysis into two SEMs to avoid losing intraspecific variability data. First, we used data at the tree level to explain RGR using the results from Objectives 1 and 2. Then, the tree-level data were summarised at the plot level to obtain a value of RGR and forest biomass per plot. These variables were then used in a subsequent SEM performed at the forest level to assess their effects on forest productivity. The model was later slightly modified as a function of the highest statistical support according to the significance of the Fisher’s C value (*P* > 0.05), indicating that the information of the missing paths does not contain meaningful information and our model fits the data (Shipley 2009). In both cases, the model was fit by maximizing the restricted log-likelihood.

Data analysis was carried out using the R statistical computing software (R Core Team, 2017). The tidyverse package was used for data manipulation and graphic display (Wickham et al., 2019). Spatial representation of the NFI was done with the package “sp” and “sf” (Pebesma, 2018). Map generation was done with the package “tmap” (Tennekes, 2018). Worldclim data was downloaded and extracted with the package “raster” (Hijmans et al., 2005). The linear mixed effect model analyses were carried out using the “lmer” package (Pinheiro et al., 2017). All the data and the script to generate these results can be accessed from an open repository in github (https://github.com/PabloCSZ/ForChange/tree/forchange_soil).

## Results

### Variation in functional traits

Functional traits showed high variability, being particularly large for leaf area (Table 1). The LD, LMA, and LT had around 14-16% coefficient of variance, and the range between minimum and maximum values was more than double. SWD and SWDMC showed the lowest coefficient of variance among all variables, ranging between 4-8% (Table 1).

The leaf nutrients were summarized through the first and second principal component axes (L_1_ and L_2_) that explained 32.7% and 20.1% of the variance, respectively (Fig. S3, Supplementary material). The axis L_1_ was mainly positively related to Cu, Fe, and Ca concentration, while the axis L_2_ was positively related to N, P, and Na concentration (Fig. S1B, Supplementary material). The intraspecific variability in functional traits related to leaf characteristics shows a strong relationship between them and was orthogonal to those related to wood density (see Fig. S2B).

The variance of plant morphological traits explained by the abiotic factors was between 0.36 to 0.61 (see R^2^ in Table 2). Aridity index was the variable with the strongest effect on LMA and leaf thickness, having a positive effect on both traits (Table 2). Abiotic factors were not able to significantly explain the intraspecific variability in LD, LA, SWDMC, and SWD in the models (Table 2). Morphological plant traits were not explained by soil nutrients (Table 2). Leaf nutrients, summarized in the first and second principal component axes (L_1_ and L_2_), were mainly explained by clay percentage (Figs. 3C and D, respectively) yet the models showed a marginal R^2^ of 0.25 and 0.02 (Table 2). A high clay percentage was related to a high Cu, Fe, and Ca leaf concentration but low N, P, and Na leaf concentration.

**Table 2.** Results of the LME models explaining the variation of functional traits with abiotic factors using a restricted maximum likelihood method of fit. Estimates for each parameter and significant correlations are indicated in bold (* *P* <0.05 and ** *P* <0.01) and nearly significant (a 0.1>*P* >0.05) in bold and italic. Marginal R^2^x100 (encompassing variance explained by only the fixed effects) and conditional R^2^x100 (comprising variance explained by both fixed and random effects are included. See Table 1 for acronyms

| Response variable | Parameters |  |  |  |  |  |
| --- | --- | --- | --- | --- | --- | --- |
| | AI | S1 | S2 | Clay % | $R^2$ M | $R^2$ C |
| LMA | **0.466\*\*** | -0.192 | 0.027 | 0.067 | 23 | 51 |
| LD | -0.04 | 0.076 | -0.139 | 0.045 | 3 | 40 |
| LT | **0.632\*\*** | -0.285a | 0.221 | 0.032 | 33 | 61 |
| LA | -0.066 | 0.25 | -0.102 | -0.248 | 4 | 41 |
| SWDMC | 0.384a | 0.035 | 0.18 | -0.073 | 8 | 35 |
| SWD | -0.145 | 0.076 | -0.087 | -0.021 | 1 | 45 |
| L1 | 0.32 | 0.143 | 0.087 | 0.293a | 25 | 79 |
| L2 | -0.046 | -0.038 | -0.097 | 0.151 | 2 | 89 |

**Figure 3.**
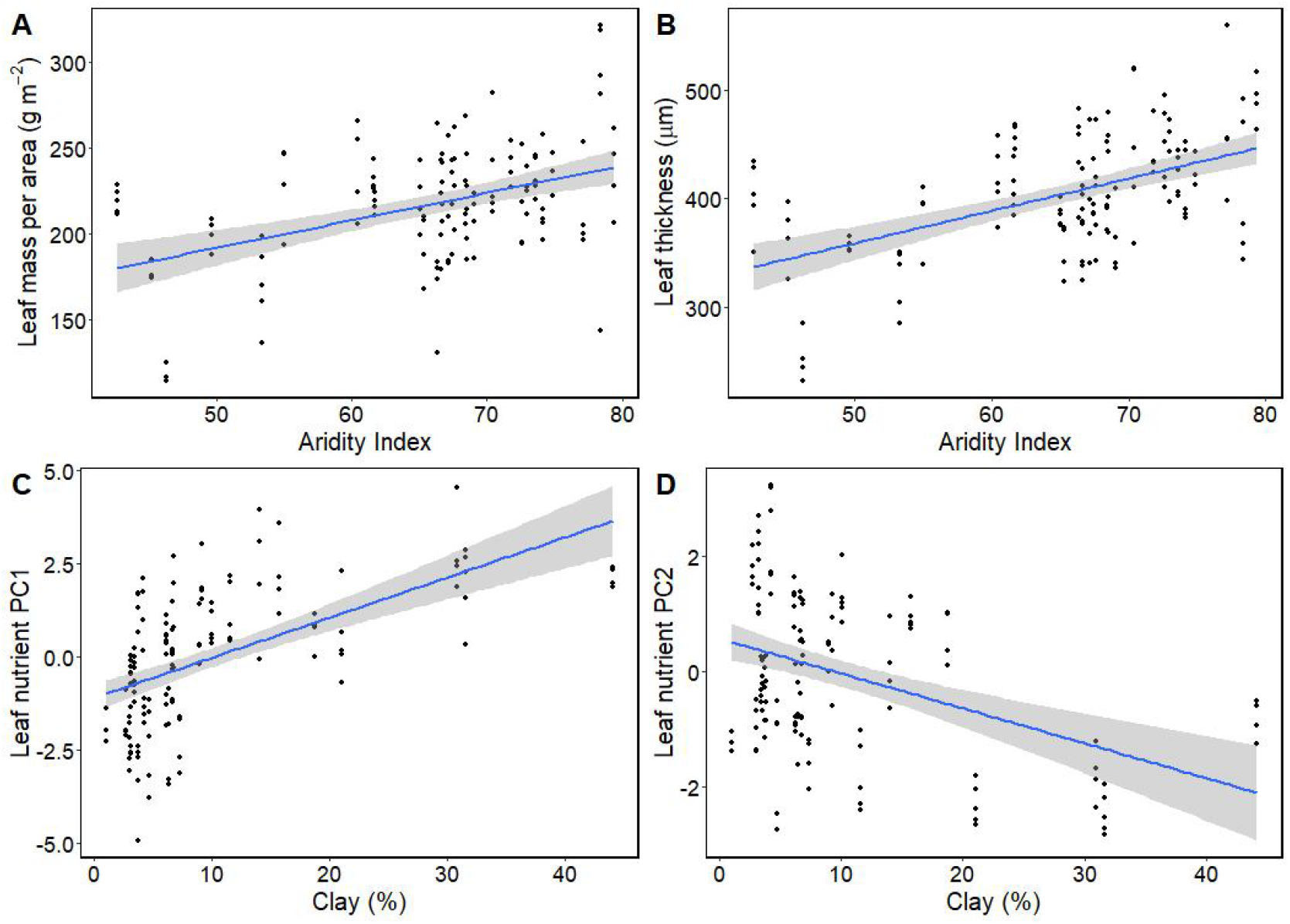
Bivariate correlation LMA and LT with Aridity (A and B, respectively), and bivariate correlation between leaf nutrient first (L_1_) and second (L_2_) PC axes with clay percentage (C and D respectively) at tree level. Grey area indicates 95% confidence interval.

### Variation in relative growth rate

The relative growth rate showed a high coefficient of variance among trees (91,74%), with values ranging between 0.45 to 236.4 g kg-1 year-1 (Table 1). It was explained negatively by tree biomass and the second axis of the soil PCA (Fig. 4A and B, respectively). Functional traits and climatic factors did not show a significant effect on RGR, and the model showed a marginal R2 of 0.27 (Table 3).

**Table 3.**
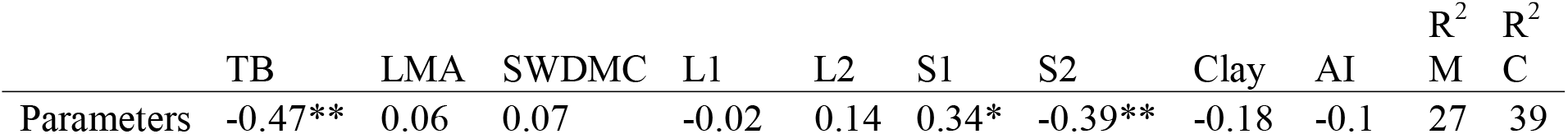
Results of the relative growth rate (RGR) model explained by tree biomass (TB), soil nutrients main PC axes (S1 and S2), soil texture, leaf functional traits (LMA, SWDMC), leaf nutrients main PC axes (L1 and L2), and aridity using a restricted maximum likelihood method of fit. Significant correlations are indicated (* *P* <0.05 and ** *P* <0.01). Marginal and conditional R^2^x100 are included.

| | TB | LMA | SWDMC | L1 | L2 | S1 | S2 | Clay | AI | $R^2$<br>M | $R^2$<br>C |
| --- | --- | --- | --- | --- | --- | --- | --- | --- | --- | --- | --- |
| Parameters | -0.47** | 0.06 | 0.07 | -0.02 | 0.14 | 0.34* | -0.39** | -0.18 | -0.1 | 27 | 39 |

**Figure 4.**
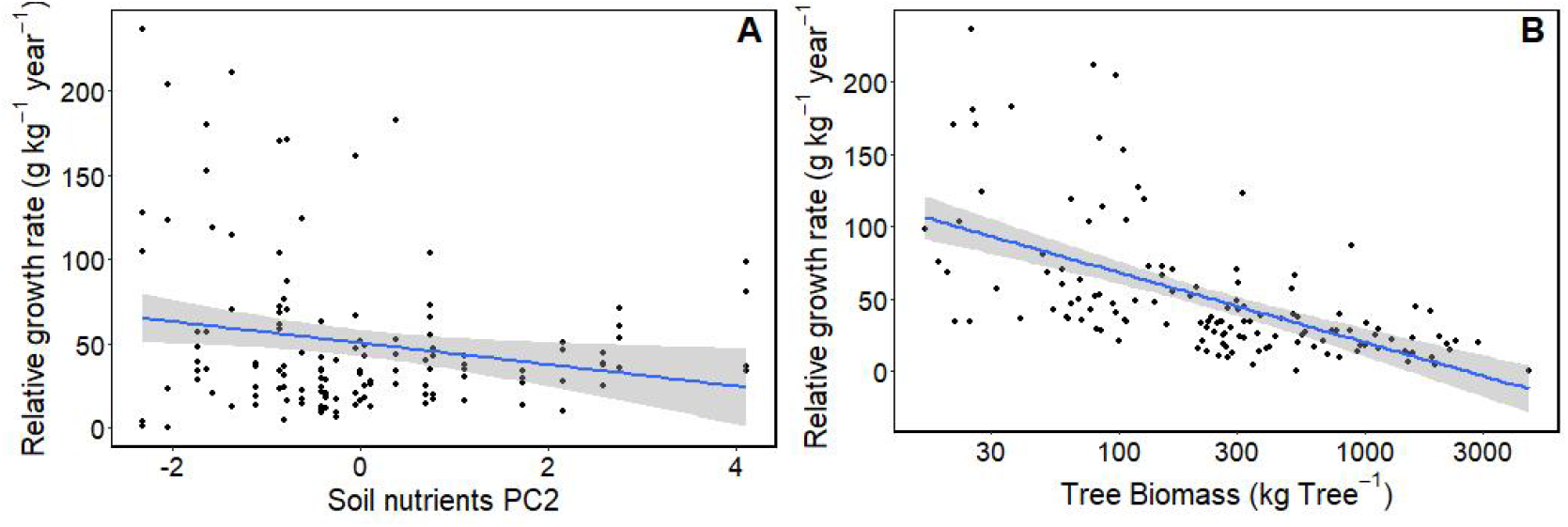
Bivariate correlation of soil nutrients PC2, and Tree biomass (axis log transformed) with the relative growth rate (A and B, respectively) at tree level. Grey area indicates 95% confidence interval.

### Forest productivity model

The two piecewise SEM built based on our initial hypothesized model showed a high goodness of fit (Fisher’s C = 6.9, *P*-value = 0.54, and Fisher’s C = 8.1, *P*-value = 0.74). At the tree level, leaf nutrients (L1) and stem-wood density explained the tree biomass and together with the soil nutrients (S1 and S2) explained the RGR of each tree. At the plot level, aridity and mean leaf nutrients (L1) explained the mean LMA of the plot, which alongside L1 explained the forest biomass. Ultimately, the mean RGR of the plot, and the forest biomass, explained the forest productivity (Fig. 5A).

**Figure 5.**
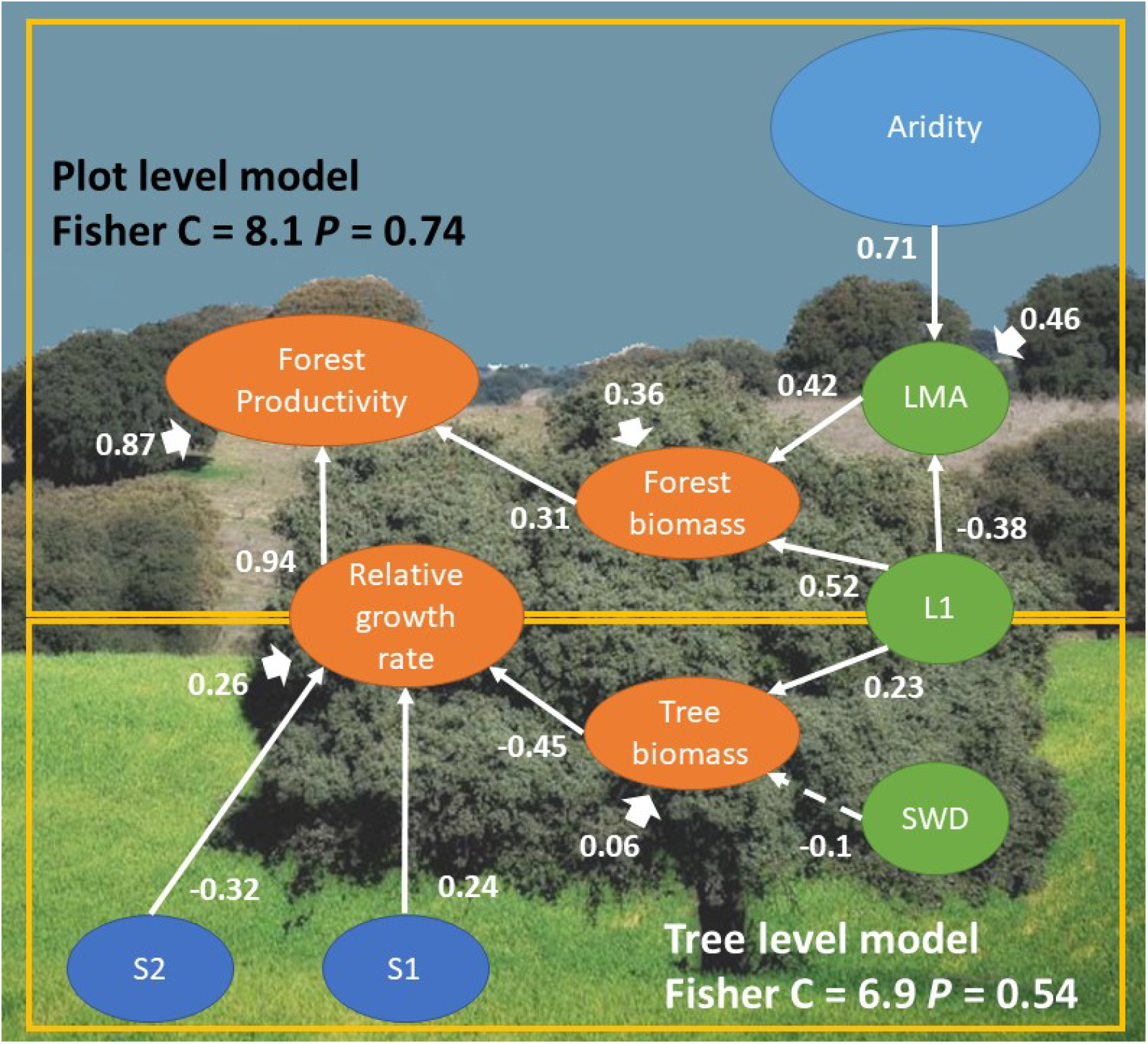
**(Top)** Piecewise structural equation modeling explaining forest productivity with forest biomass and RGR based on our initial hypothesis (see Fig. 1) using data from 30 plots. **(Bottom)** Piecewise structural equation modeling explaining the RGR of each tree in our study (n = 135). Dashed lines indicate non significant relationships. Short arrows indicate the correlation coefficient for each response variable (n = 134). S1 and S2 (first and second axes of soil PCA), LMA (leaf mass per area).

## Discussion

Our results showed that forest productivity was mainly explained by RGR, which, in turn, was affected by tree biomass and soil nutrients, with an indirect effect of functional traits and climate factors. We found a positive effect of LMA and leaf nutrients on forest biomass that directly affected forest productivity. This suggests that the variability of functional traits in response to climate factors might directly affect forest carbon storage in the long run.

### Effects of abiotic factors on functional traits

LMA constitutes the most measured and commonly used trait to describe plant ecological strategies. Our results indicated that LMA was positively related to aridity. Similarly, it has also been found to decrease with precipitation (Ogaya and Peñuelas, 2007) and temperature (Poorter et al., 2009; Vilà-Cabrera et al., 2015). Leaf thickness was also explained by aridity and showed a high correlation with LMA. This suggests that the increase in LMA under drought conditions is linked with an increase in leaf thickness, associated with a larger volume of mesophyll area in the leaf tissue (Villar et al., 2013). In theory, this morphological adaptation has a direct impact on the gas exchange (Maire et al., 2015) and water transport (Rosas et al., 2019), and it may reduce the impact of abiotic stress on plant growth.

The leaf chemical traits were mainly explained by the clay percentage, rather than aridity or soil nutrients. A high clay percentage in the soil is generally associated with low water drainage and high nutrient storage (Anderson et al., 2006). Overall, this leads to a high amount of soil nutrients (Table S1, Supplementary material), and ultimately, a high concentration of nutrients in the leaf. However, for some nutrients, a high clay percentage associated with a high concentration of iron oxide can also lead to a more negative soil matric potential in summer and autumn and reduce soil nutrient transport, a situation referred to as the inverse texture effect (Fernandez-Illescas et al., 2001). This might be our case based on the negative correlation between clay percentage and the PC2 leaf axis (mainly explained by the leaf N and P concentration), and the negative effect of clay% in soil P (data not shown). Furthermore, an increase in leaf phosphorus concentration seems to be paired up with high leaf manganese concentration (Fig. S1). The accumulation of Mn in the leaf is relatively common when P mobility from the soil is limited (Lambers et al., 2015). It seems that root exudates can locally modify the pH, which makes P more available alongside Mn (White et al., 2013). Therefore, it is possible that the clay percentage in the soil can increase soil nutrients concentration, but reducing its mobility in highly demanded nutrients, like phosphorus.

### Effect of abiotic factors, functional traits, and forest structure on RGR

The individual tree biomass was the main intrinsic factor explaining the RGR of the trees. This negative relationship suggests larger trees tend to grow more slowly, which could be explained by a lower proportion of leaves (Villar et al., 2017). Thus, as the tree biomass and the number of resources required for growing increase, nutrient assimilation and transport proportionally decrease, reducing RGR. Competitive interactions for light and nutrients between surrounding trees reduce plant growth, at least in the early years (Binkley, 2004). Thus, the magnitude of the biomass-growth relationship depends on the competitive environment (Ruiz-Benito et al., 2015). Furthermore, hydraulic and nutrient limitations appear in older trees due to mechanical constraints in vessel size, which reduces both water and nutrient transport (Mencuccini et al., 2005). An increase in soil nutrient concentration should yield high plant biomass, however, this only seems to be true for small trees (Li et al., 2018). The increase in soil nutrients concentration does not mitigate the morphological changes associated with nutrient transport when the tree gets older and the biomass increases (Drake et al., 2010). This could explain the lack of relationship between RGR and the soil nutrients summarized in the first PCA axis, which is mainly explained by Ca and N content in the soil. However, opposite results have also been found (Paoli and Curran, 2007; Zemunik et al., 2018), and further studies are still required to understand the effect of soil nutrients on plant growth.

Climate factors had a low effect on RGR, in agreement with previous results for several *Quercus* species suggesting a lower explanatory power of climate in comparison to local factors as nutrient availability, soil texture, and tree density (Villar et al., 2017). Furthermore, functional traits did not show a significant effect on RGR either, which contradicts several studies that suggested functional trait variability has a physiological impact at the whole-plant level (Poorter et al., 2009; Violle et al., 2007). However, most of those studies have been performed for young seedlings rather than adult trees (Bastias et al., 2018). When this distinction has been made, results have shown that LMA does not correlate with RGR for adult trees (Laughlin et al., 2017; Wright et al., 2010). The lack of relationship has been justified by the absence of a clear physiological basis in the so-called “soft traits’’, like LMA and leaf thickness, and tree height (Rosas et al., 2019). However, this kind of terminology has been questioned because it focuses on operational data measurements (how easy or difficult it is to measure a certain trait) and underestimates the functional meaning of leaf morphology (Violle et al., 2007). In *Q. ilex* evergreen leaves, high LMA is the result of high sclerenchymatic tissues that confer leaf resistance to water diffusion from the vein to the mesophyll during climatic stress (Enrique G. de la Riva et al., 2016). Most likely, the leaf functional trait plasticity is reducing the negative effect of environmental stress, thus, limiting the reduction on plant fitness in RGR (Gratani, 2014). According to this, morphological adaptations in the leaf are made to keep plant growth steady regardless of climate conditions.

### Modeling forest productivity depending on ecological, physiological, and environmental factors

Our models confirm our initial hypothesis, which suggests that forest productivity depends on both the vegetation quality (RGR) and the quantity (forest biomass). Key functional traits like LMA and SWD are affected by forest productivity, but not directly as we initially hypothesized. For instance, the effect of LMA and climatic factors (i.e. aridity) over RGR did not occur directly, instead, increases of leaf area relative to leaf weight directly affected tree growth rates (Antúnez et al., 2001; Lambers and Poorter, 2004). However, this relationship does not hold itself when the tree size increases because respiratory losses for maintenance increases with the mass in large trees (Mencuccini et al., 2005). Thus, even though LMA is a highly sensitive trait to environmental factors, its variation does not always represent an adaptive advantage to increase the RGR. Instead, we found a positive effect of LMA on forest biomass. This suggests that a thick and dense leaf morphology increases forest biomass in the long term. Similar results have been found when limited conditions are present (van der Sande et al., 2018), and in temperate forests (Yuan et al., 2019). Therefore, leaf morpho-physiological changes might be an adaptive strategy to maintain water transport and C uptake under abiotic stress, therefore increasing the forest biomass.

Ultimately, our study showed forest productivity is mainly explained by RGR, which is affected by tree biomass and soil nutrients. This result suggests that protecting established forest populations increases forest biomass, and utterly, forest productivity (Morán-Ordóñez et al., 2020). Scheduling pruning and wood harvesting allow for a reduction in stand density, increase light availability in dense patches, and eliminate competitive interactions between trees (Reyes et al., 2008). Similarly, protecting species diversity and regulating animal foraging increase soil fertility in the long term (Ammer, 2019; James et al., 2009). Thus, the implementation of regional forest policies may be as important as national environmental regulations to protect *Q. ilex* forest productivity (Mokany et al., 2008).

## Conclusion

Our findings highlight the large intraspecific variation in key functional traits (i.e. LMA) were associated with a climatic gradient and shows that plant adaptations to abiotic stress do not always have direct effects on plant growth or forest productivity, but indirect effects can be important. Instead, soil fertility and tree biomass have a fundamental effect on the relative growth rate, suggesting forest management may have a bigger role than we originally thought to maintain forest ecosystem services.

## Acknowledgements

Financial support was provided by the project *Ecología funcional de los bosques andaluces y predicciones sobre sus cambios futuros* (For-Change) (UCO-27943) from Junta de Andalucía (Spain) and by the Spanish MEC project DIVERBOS (CGL2011-30285-C02-02) and ECO-MEDIT (CGL2014-53236-R) and FEDER funds. We thank the MAPA (Ministerio de Agricultura, Pesca y Alimentación) and MITECO (Ministerio de Transición Ecológica) the access and the open-access availability of the Spanish Forest Inventory (https://www.mapa.gob.es/). CCB was supported by a postdoctoral fellowship of the Ramón Areces Foundation.

## Notes

https://github.com/PabloCSZ/ForChange/tree/forchange_soil

### Competing Interest Statement

The authors have declared no competing interest.

