## Supplementary Table and figures for "Linking functional traits with tree growth and forest productivity in *Quercus ilex* forests along a climatic gradient"

**Figure S1. (Top)** Principal component analysis (PCA) of leaf nutrients in *Quercus ilex* studied plots. Normal confidence ellipses (95% interval) for each province is shown. **(Bottom)** Variable scores for the principal component analysis.

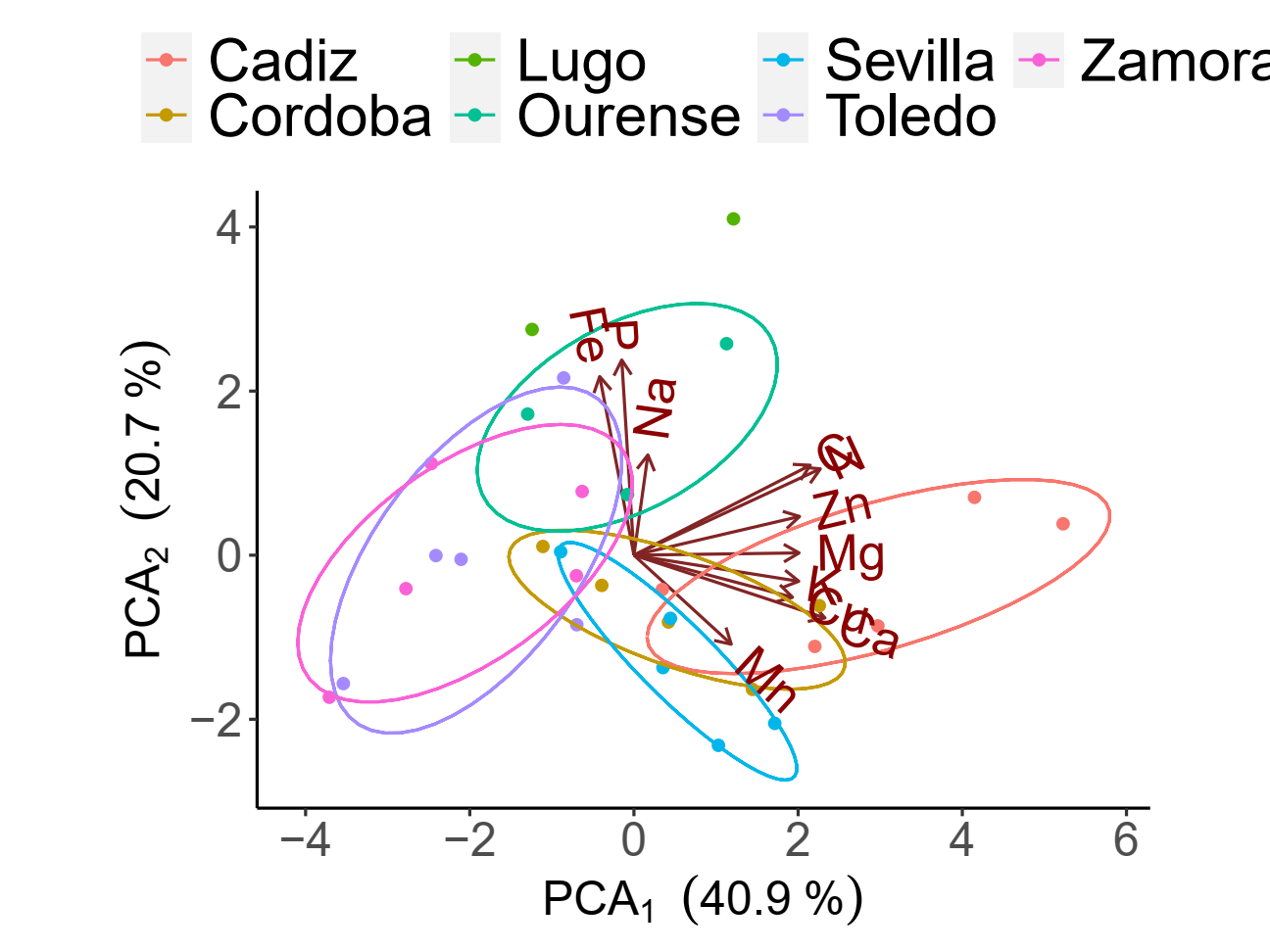

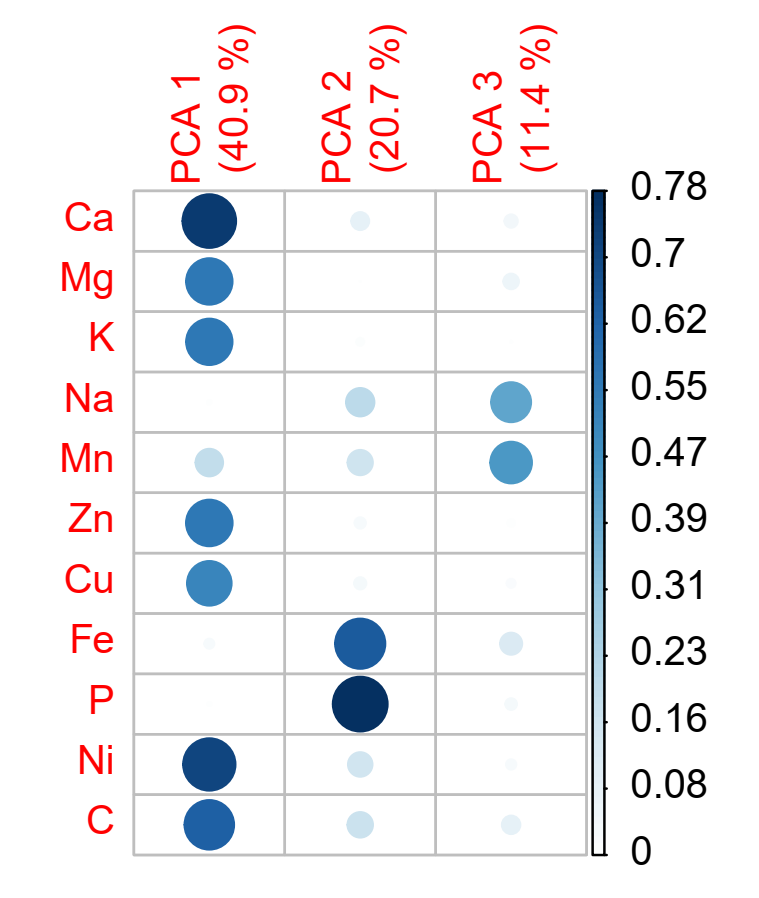

**Figure S2.** **(Top)** Bivariate correlation matrix between abiotic factors. Soil nutrient PCA axes (S1 and S2), MAT: mean annual temperature, MAP: mean annual precipitation, and AI: aridity. **(Bottom)** Bivariate correlation matrix between functional traits. LT: leaf thickness, LMA: leaf mass per area, LD: leaf density, LA: leaf area, and SWDMC: stem-wood dry matter content, and SWD: stem-wood density.

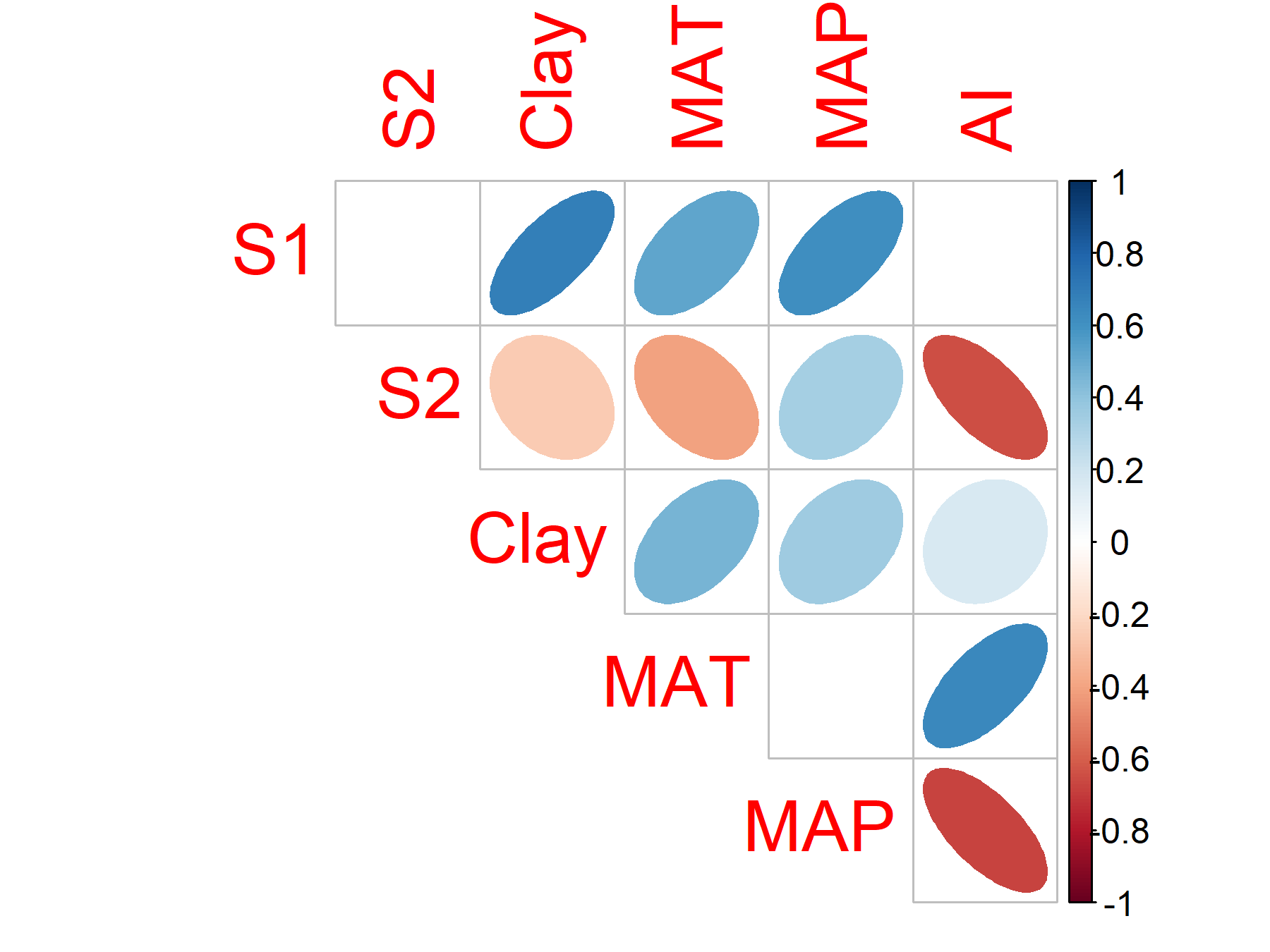

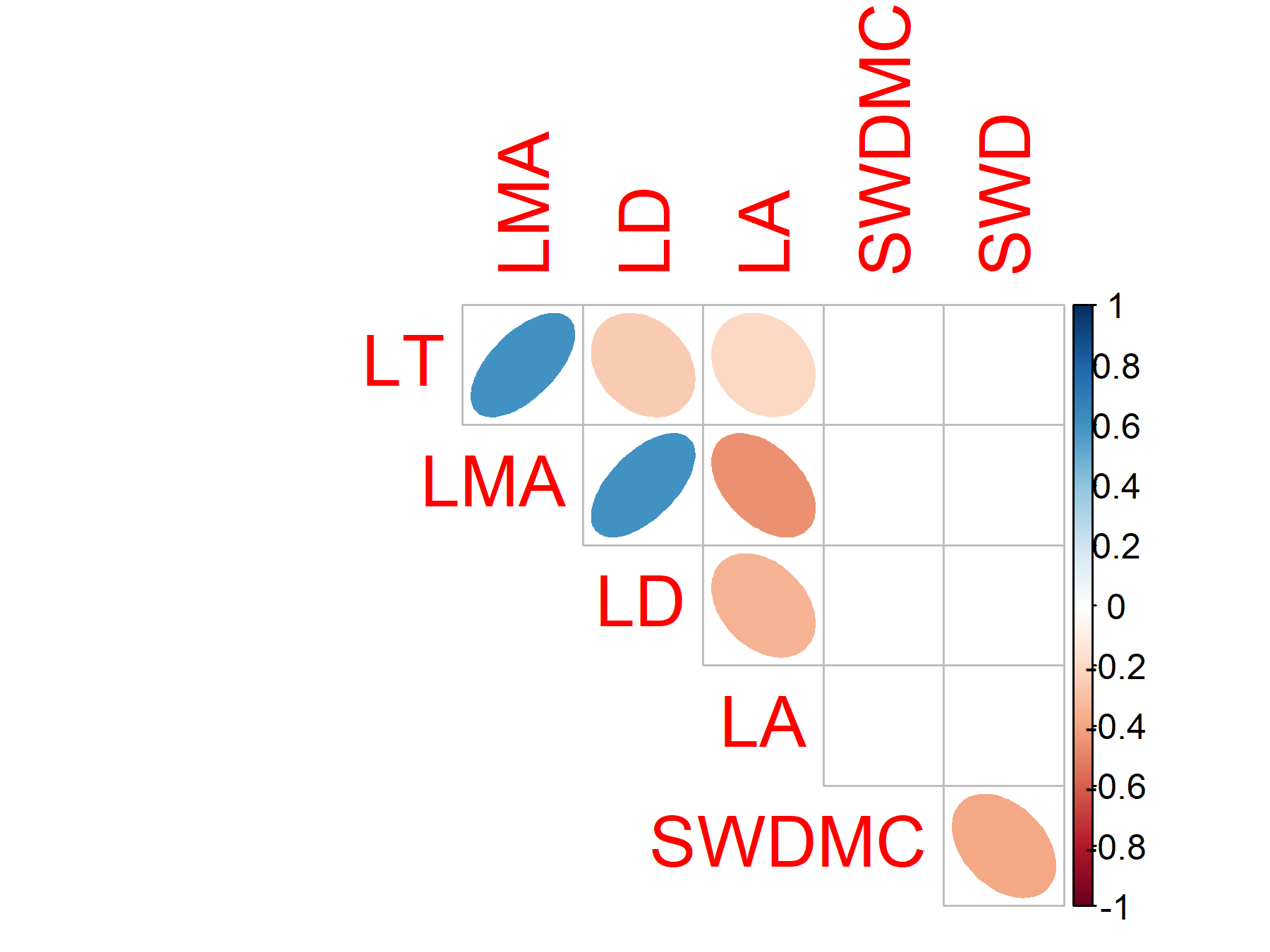

**Figure S3. (Top)** Principal component analysis (PCA) of soil nutrients. Normal confidence ellipses (95% interval) for each province is shown. **(Bottom)** Variable scores for the principal component analysis.

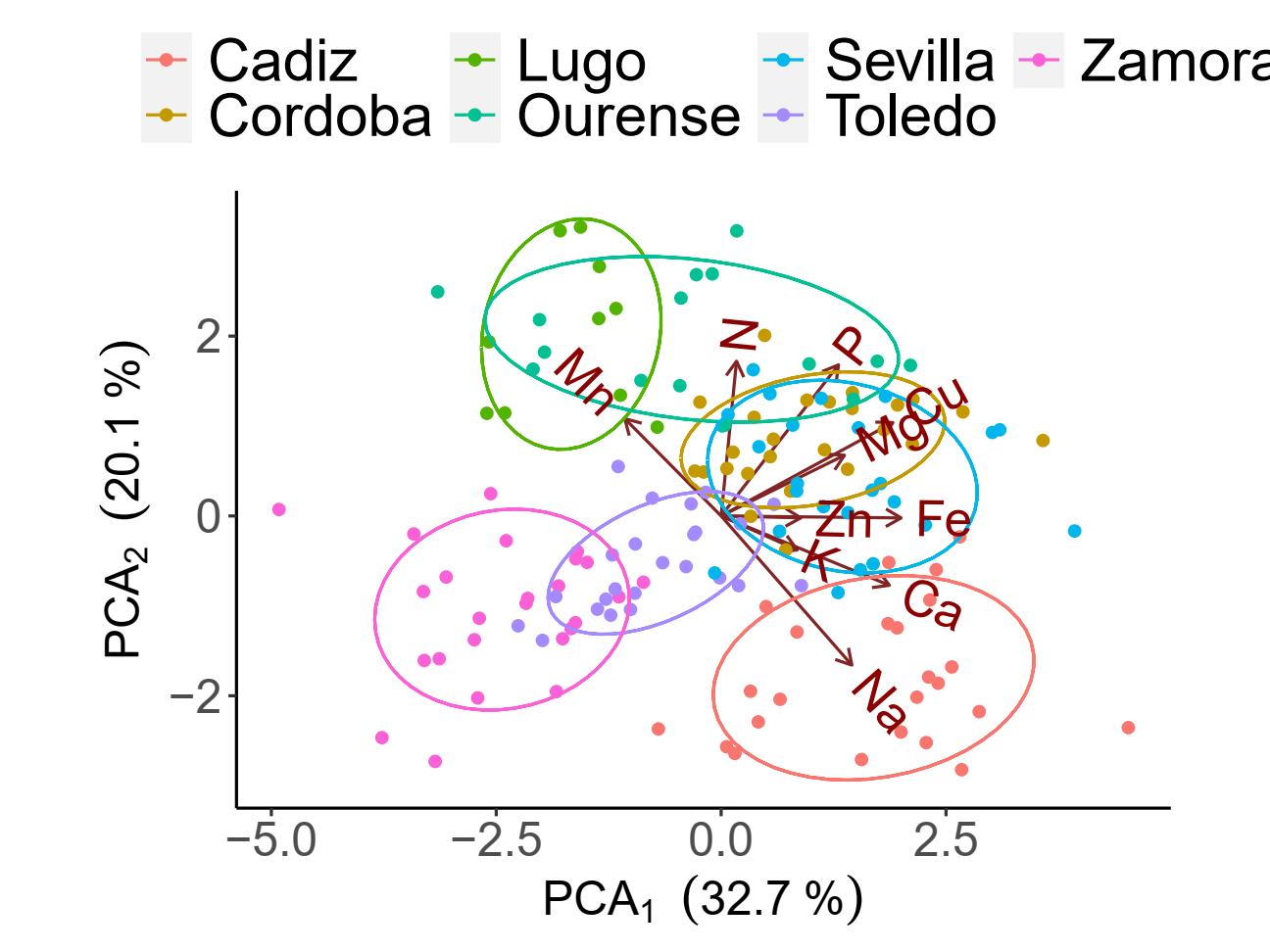

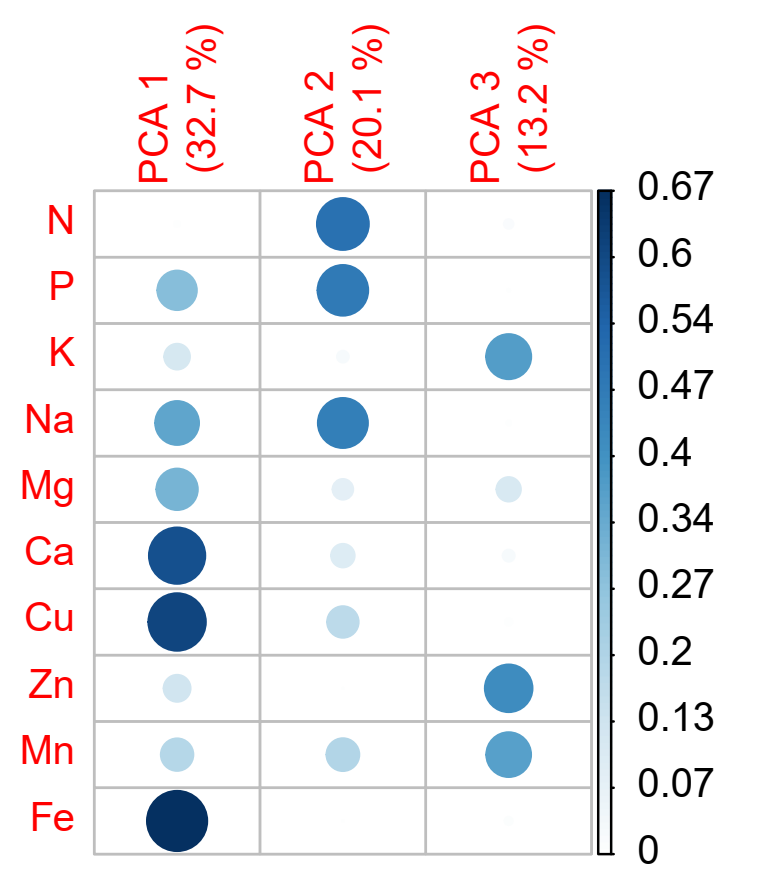

**Table S1.** Descriptive statistics of soil and leaf nutrients.

|  | **variable** | **mean** | **sd** | **min** | **max** | **cv** |
| --- | --- | --- | --- | --- | --- | --- |
| Soil | C | 3.54 | 2.92 | 0.84 | 12.01 | 82.35 |
|  | Ca | 2328 | 4226 | 81 | 14712 | 181.5 |
|  | Cu | 3.29 | 2.10 | 1.31 | 11.19 | 63.69 |
|  | Fe | 58.67 | 72.66 | 12.64 | 349.01 | 123.8 |
|  | K | 126 | 61 | 65 | 360 | 48.70 |
|  | Mg | 254 | 162 | 52 | 758 | 63.99 |
|  | Mn | 63.29 | 46.86 | 16.88 | 248.15 | 74.04 |
|  | Na | 301 | 64 | 231 | 499 | 21.40 |
|  | Ni | 0.19 | 0.11 | 0.06 | 0.59 | 57.43 |
|  | P | 16.57 | 20.40 | 2.81 | 108.47 | 123.1 |
|  | Zn | 1.10 | 1.26 | 0.07 | 4.23 | 114.8 |
| leaf | C | 47.77 | 1.05 | 45.22 | 50.39 | 2.19 |
|  | Ca | 5232 | 2385 | 1141 | 12956 | 45.59 |
|  | Cu | 3.76 | 1.40 | 1.44 | 8.37 | 37.16 |
|  | Fe | 248 | 243 | 18 | 1433 | 97.97 |
|  | K | 4573 | 1172 | 2353 | 8170 | 25.62 |
|  | Mg | 1607 | 598 | 525 | 3383 | 37.19 |
|  | Mn | 1324 | 873 | 70 | 4239 | 65.89 |
|  | N | 1.29 | 0.27 | 0.67 | 2.29 | 21.08 |
|  | Na | 133 | 82 | 31 | 677 | 61.48 |
|  | P | 503 | 298 | 97 | 1323 | 59.19 |
|  | Zn | 23.81 | 7.44 | 10.95 | 48.34 | 31.26 |
